# Multiomic Integration Reveals Taxonomic Shifts Correlate to Serum Cytokines in an Antibiotics Model of Gut Microbiome Disruption

**DOI:** 10.1101/2025.02.15.638396

**Authors:** Cameron X. Villarreal, Deva D. Chan

**Author notes:** Corresponding Author: Deva Chan.

## Abstract

The gut microbiome interacts with many systems throughout the human body. Microbiome disruption reduces bone tissue mechanics but paradoxically slows osteoarthritis progression. The microbiome also mediates inflammatory and immune responses, including serum cytokines. Towards our long-term goal of studying how the gut microbiome interacts with synovial joint health and disease, we examined how antibiotics-induced changes to microbial taxa abundance associated to serum cytokine levels.

Mice (n = 5+) were provided ad libitum access to water containing antibiotics (1 g/L neomycin, 1 g/L ampicillin, or 1 g/L ampicillin with 0.5 g/L neomycin) or control water from 5- to 16-weeks old, corresponding in skeletal development to ∼10 to ∼25 years in humans. At humane euthanasia, we collected cecum contents for 16S metagenomics and blood for serum cytokine quantification for comparison to control and among antibiotic groups. We used dimensional reduction techniques, multiomic integration, and correlation to discriminate antibiotic groups and identify specific relationships between high-abundance taxa and serum cytokines.

Antibiotic treatment significantly lowered diversity, altered phylum relative abundance, and resulted in significant association with specific taxa. Dimensional reduction techniques and multiomic integration revealed distinct antibiotic-associated clusters based on genera relative abundance and cytokine serum concentration. Cytokines IL-6, MIP-1B, and IL-10 significantly contributed to antibiotic discrimination, significantly different among antibiotic treatments, and had significant correlations with specific taxa.

Antibiotic treatment resulted in heterogenous response in gut microbiome and serum cytokines, allowing significant microbe-cytokine links to emerge. The relationships identified here will enable further investigation of the gut microbiome’s role in modifying joint health and disease.

## INTRODUCTION

The gut microbiome is a vast community of microbes that resides in the intestinal tract. Characteristics of the gut microbiome, including the abundance of constitutive taxa and the diversity of those taxa, have been connected to many physiologic and pathophysiologic systems, including maintenance and disease in synovial joints. The gut microbiome can modulate nutrient metabolism and extraction [1], vitamin metabolism [2], and host inflammatory response [3]. Beyond these indirect associations, the gut microbiome mediates rheumatoid arthritis pathogenesis through host immune response modulation [4]. Additionally, the gut microbiome was markedly disrupted in osteoarthritis patients [5], and probiotic supplementation protected against postmenopausal bone loss [6]. These associations suggest that the gut microbiome may modulate synovial joint health and disease. However, our limited knowledge of the potential mechanisms through which this crosstalk occurs hinders our ability to treat and study gut-joint-related diseases.

Congruent to findings in human subjects, a wealth of evidence has shown that the gut microbiome plays a role in both musculoskeletal health and disease in animal models [7–9]. Recent work has shown that microbiome manipulation has been shown to result in reduced bone tissue strength and crystallinity [10, 11]. Beyond tissue level strength, gut microbiome manipulation may alter the incidence, progression, and severity of joint degradation. A high-fat diet has been shown to cause taxonomic shifts in the microbiome and promote joint degradation in an osteoarthritis model in rats [12]. Additionally, in a model of post-traumatic osteoarthritis, germ-free mice had higher bone volume fraction and trabecular thickness following ACL rupture compared to conventional mice [13]. While these links exist in humans and animals, unknown systemic mediators that may link the gut microbiome to synovial joint health and disease remain poorly understood.

Inflammation is a potential mediator through which the gut microbiome may alter joint health and disease. Microbial community diversity [14] and relative abundance of constitutive taxa within the community [15] are associated with metabolic syndromes and other inflammatory diseases comorbid with musculoskeletal disease. For example, taxonomic shifts are indicative of obesity-related inflammation [16, 17] and type II diabetes [18, 19], both of which are comorbid or associated with joint diseases [20, 21]. Furthermore, patients with disrupted gut microbiota evidenced by reduction or increase of specific taxa have displayed concomitant alteration in inflammatory signaling in a variety of diseases [22–24]. Thus, alterations to the microbial community typically result in altered inflammation, although it is difficult to ascertain the direction of this interaction in a clinical setting.

Similarly, the gut microbiome alters systemic inflammation in animal models. Recent work has shown that gut microbiome manipulation through antibiotic use alters systemic inflammation in mice, evidenced by serum cytokine concentration levels in mice [25, 26]. However, these studies examine short timespans (only up to 4 weeks) and therefore examine a more acute response to antibiotics administration during skeletal maturation. Furthermore, gut microbiome homogenization to a stimulus typically occurs within social housing of mice on a similar timeline (3-4 weeks) [27, 28]. Therefore, extending this timeline past when the gut microbiome tends to homogenize and up to skeletal maturity may provide deeper insight into microbiome driven systemic effects after antibiotic treatment and allow a deeper characterization of the circulating cytokine profile during a longer period of skeletal maturation [29, 30].

Although it remains unclear how microbiome manipulation may drive systemic inflammation. The connection between inflammatory signaling and synovial joint health is well documented. For example, IL-6 situationally regulates both bone resorption and restitution [31, 32], IL-10 improves bone restitution [33], and MIP-1β increases osteoclast differentiation in multiple myeloma [34]. Additionally, IL-6 [35] is associated with osteoarthritis related cartilage loss in and MIP-1β and IL-10 concentrations are increased in the synovial fluid of osteoarthritis patients [36]. IL-6 [37, 38] and IL-10 [39] also regulate tendon and ligament turnover. Furthermore, IL-6 regulates catabolic protein expression in meniscus cells [40], adipocyte function [41], and, in conjunction with IL-10, shapes the bone marrow microenvironment and cell differentiation therein [42, 43]. The gut microbiome’s ability to affect changes in cytokine secretion and the role of cytokines in musculoskeletal tissue maintenance therefore suggest a potential mechanism through which the gut microbiome could alter joint health and disease.

Towards our long-term goal of examining potential mechanisms through which the gut microbiome may modulate musculoskeletal health, we first aimed to characterize the links between the microbiome and systemic inflammatory cytokine changes in antibiotics-induced models of gut microbiome disruption. In this study, our objectives were to determine how, during skeletal maturation in mice, (1) different antibiotic treatments alter the gut microbiome composition, (2) antibiotic use alters systemic inflammation, (3) and antibiotic-associated gut microbiome and systemic inflammation changes relate. We hypothesized that both gut microbiome population and systemic inflammation would be altered in an antibiotic-specific manner. We tested this by treating mice with ampicillin, neomycin, or a cocktail of ampicillin and neomycin from before sexual maturity (5 weeks) to skeletal maturity (16 weeks) and quantifying changes to gut microbiome taxonomy and serum cytokine concentration. This time frame also enables study of circulating cytokines during skeletal maturation, a time during which signaling molecules play a key role in the maintenance and growth of musculoskeletal tissue [44, 45]. Additionally, using differing antibiotic treatments enables study of the heterogeneity in microbiome disruption [11], and therefore potential heterogeneity in downstream effects on systemic cytokine concentration. To examine these relationships, we applied dimensional reduction, multiomic integration, and univariate statistical methods to quantify the association between gut microbiome manipulation and altered systemic inflammation in mice treated with antibiotics up to skeletal maturity. To the best of our knowledge this is the first time multiomic integration has been used to examine specific links between the constituent taxa in the gut microbiome and circulating cytokine concentrations resulting from antibiotic disruption. Understanding the antibiotic-specific changes to the gut microbiome and systemic cytokines through this combination of statistical methods would then enable the identification of potential microbial taxa and cytokine pathway targets to study the gut-joint axis.

## METHODS

### Animal Experiments

No more than 5 mice per cage were housed on ventilated racks, provided *ad libitum* access to conventional laboratory diet (chow) and sterilized drinking water, and exposed to standard 12-hour light-dark cycles. Mice were randomly assigned by cage (at least two cages per group) to control (8 male (M)/8 female (F)) or antibiotics-treated groups. 1 g/L ampicillin (Amp, 6F/4M), 1 g/L neomycin (Neo, 5M/5F), or 1 g/L ampicillin + 0.5 g/L neomycin (Amp+Neo, 8M/8F) was added to drinking water with no change to chow, from 5 to 16 weeks of age. Minimum group sizes for cytokine analysis were determined via prior precedent [46], and verified via power analysis. We selected the concentration of the Amp+Neo combination to replicate an established model that results in gut microbiome disruption and downstream physiologic effects and ensures mice will consume the treated water without need for added sweeteners [47, 48]. To determine treatment duration effects, we treated an additional set of mice from 5 to 8 weeks of age (6 M, 6 F) with Amp+Neo – a short-term treatment model – and compared against age-matched controls. Control mice were offered normal water and chow. During treatment, no adverse health outcomes were observed, and thus no animals were excluded. At euthanasia, whole blood was collected, allowed to coagulate on ice, and then centrifuged at 1500*g* for 10 minutes at 4°C. Serum was collected and stored at −80°C until analysis. Cecum contents were collected under sterile conditions and stored in DNA/RNA Shield stabilization reagent at 4°C (Zymo Research Corp., Irvine, CA) until samples were shipped for sequencing.

### Gut Microbiome Taxonomic Characterization and Analysis

The gut microbiota composition was quantified via 16S rRNA sequencing using commercial services (Zymo Research Corp., Irvine, CA). The V3-V4 region of the 16S rRNA gene was amplified and sequenced on Illumina MiSeq with a v3 reagent kit (600 cycles). Unique amplicon sequences (single DNA sequences) were inferred from raw reads using the DADA2 pipeline [49], and taxonomy was assigned using Zymo’s established pipeline. Relative abundance was defined as the relative proportion of a specific taxa relative to the total community. Shannon diversity was used as the alpha diversity metric since it does not weight rare species which may be less susceptible to antibiotics [50].

To identify microbial community similarity and difference, we first evaluated phylum-level relative abundances and the overall microbiome diversity. The microbiome composition of mice and humans share marked similarity, with the phyla Firmicutes, Bacteroidetes, Actinobacteria, Proteobacteria, Tenericutes, and Verrucomicrobia being among the most abundant in both human and mouse [51, 52]. Shannon diversity [53], *H*, quantifies the diversity and evenness of a community [54] and is calculated as 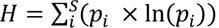, where *p_i_* is the proportion of the *i*^th^ species and *S* is the number of species present. We compared phylum-level relative abundances and Shannon diversity among antibiotics-treated and control groups using Kruskal-Wallis test with post hoc pairwise comparisons to control via Dunn test with Benjamini-Hochberg correction for multiple comparisons. We also compared Shannon diversity in short-term Amp+Neo treatment duration to age-matched controls using Mann-Whitney U test against control. Non-parametric tests were selected to handle non-normally distributed data (assessed via Shapiro-Wilk test) or bounded data set. Statistical testing was performed in RStudio [55], running R (version 4.4.2) and using the rstatix (version 0.7.2) package. Unless otherwise stated, α = 0.05 was defined as a threshold for statistical significance.

To evaluate the microbiome at a more granular scale, we next plotted unique and shared amplicon sequence variants (ASVs), among treatment groups filtered to have at least 50 reads to mitigate sampling error, in a Venn diagram [56]. We identified the top 30 conserved genera among all groups and performed principal component analysis to observe clustering due to the relative abundance of these genera. To identify significant associations between treatment and taxonomic shifts at the genus level relative to control, we conducted Microbiome Multivariate Associations with Linear Models (MaAsLin2) [57] with a minimum prevalence of taxa set at 5%, treatment as the fixed effect, and post hoc q values calculated using the Benjamini-Hotchberg method.

### Serum Cytokine Concentration Quantification and Analysis

To measure circulating cytokines in serum, magnetic bead assays were performed using a MILLIPLEX mouse 32-plex cytokine/chemokine magnetic bead panel (Millipore, MCYTOMAG-70K-PX32), adapted for 384-well plates and small sample volumes [46, 58]. Samples, blanks, and quality control were loaded in triplicate, and cytokine concentrations were quantified with the Luminex Bio-Plex 3D system. Cytokines with 25% or more of the samples from any single group reading below the lower limit of quantification were excluded from comparison to control for that group. These thresholds have been verified in other published work [46]. The cytokines that exceeded this threshold were MIP-1a, IL-4, IL-12 p70, IL-15, and IL-17 and GM-CSF, leaving 26 remaining cytokines. As with phyla, we compared cytokine concentrations among treatment groups using Kruskal-Wallis test, followed by post hoc Dunn test with Benjamini-Hochberg correction. We then used the Spearman correlation to identify significant correlations and linear trends between 16S taxonomic data and serum cytokine concentrations.

To identify the magnitude and significance of differences in the inflammatory profile of antibiotics treatment groups relative to control at long- and short-term treatment duration, we created volcano plots of the serum cytokine log_2_(fold change) vs. – log_10_(*p*), where p is the p-value from Mann-Whitney U test against control. We defined fold change as 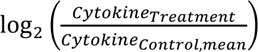. To capture subtle but meaningful differences in serum cytokine concentrations, we used a cutoff of log_2_(fold change) > 0.5, which has been used in other contexts [59]. We used Kruskal-Wallis, adjusted for false discovery rate (FDR), followed by pairwise Mann-Whitney U test with post hoc Benjamini-Hotchberg test for all samples meeting the threshold q < 0.1 for FDR.

### Supervised Dimensional Reduction and Multiomic Integration

To determine discriminating cytokine signatures among antibiotic treatments and to examine the effect of treatment duration in Amp+Neo mice, we implemented sparse partial least squares discriminant analysis (sPLS-DA), a supervised dimensional reduction technique, using the mixOmics package [60]. To determine how taxonomy and inflammatory signaling together discriminate among antibiotic treatments, we performed sPLS-DA with genera relative abundance and fold change serum cytokine concentration. We determined the variables most contributing to group separation via variable importance projection (VIP), where a VIP > 1 indicates that this variable is among the most relevant in explaining the separation between groups. We next performed Data Integration Analysis for Biomarker discovery using Latent cOmponents (DIABLO) [61] to identify related features within the 16S and cytokine datasets, determine correlations between datasets, and optimize discrimination among treatment groups. We assessed the performance of the DIABLO model via centroid distance weighted error rate and area under the curve.

## RESULTS

### Antibiotics Treatment Results in Broad Changes to the Gut Microbiome

We compared the relative abundance of specific taxa at the Phylum level (Figure 1a) to characterize broad changes to the gut microbiome between antibiotic treatments and control and among antibiotic treatments. The relative abundance of Tenericutes (p = 0.015) was significantly lower while that of Verrucomicrobia (p = 0.002) was significantly higher in Amp compared to control. In Neo, the relative abundance of Tenericutes (p = 0.002) and Actinobacteria (p = 0.014) was significantly lower than that of control. In Amp+Neo the relative abundance of Bacteroidetes (p = 0.040) and Proteobacteria (p = 0.008) were significantly lower than control, and the relative abundance of Actinobacteria (p <0.001) and Verrucomicrobia (p < 0.001) were significantly higher than control. Shannon diversity was significantly lower (p < 0.001) than control in all antibiotic treatment groups (Figure 1b). The divergent relative abundance of these phyla among antibiotic groups demonstrate that the microbial community has been disrupted in an antibiotic specific manner.

**Figure 1.**
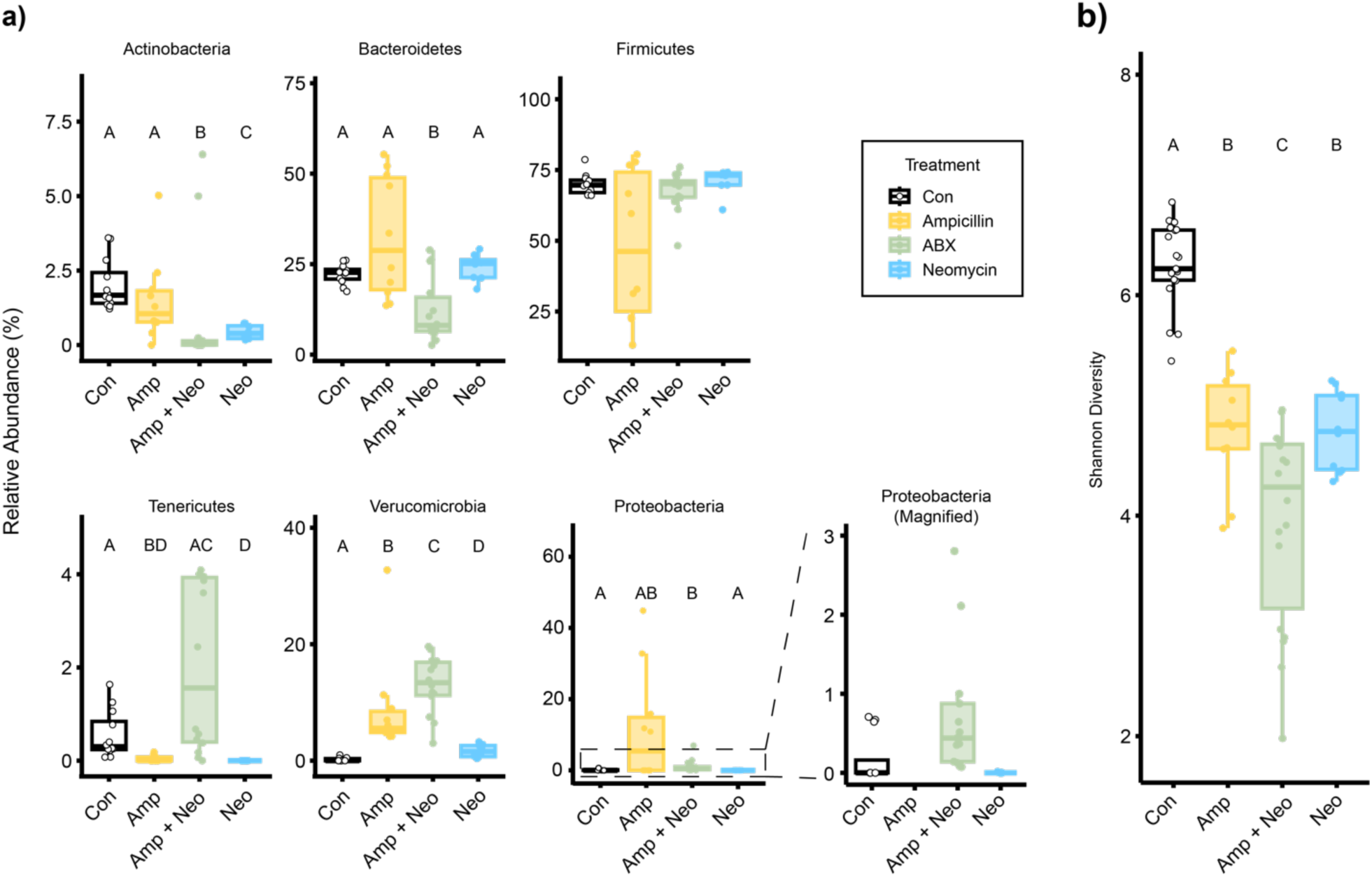
Antibiotic treatments result in heterogeneous modification to the gut microbiome. The gut microbial communities in control (Con) and 1 g/L ampicillin (Amp), 1 g/L neomycin (Neo), or 1 g/L ampicillin + 0.5 g/L neomycin (Amp+Neo) treated mice were characterized using (a) relative abundance of phyla and (b) Shannon diversity. Amp data is removed from the Proteobacteria magnified inset for clarity. Treatments were compared via Kruskal Wallis, with significant effects evaluated by a post hoc Dunn test with Benjamini-Hochberg correction. Treatment groups with unique letters are significantly different from each other (p<0.05).

### Granular Separation of the Gut Microbiome Leads to Separation between Treatment Groups

We first examined community similarity by comparing the number of conserved and unique ASVs among treatment groups and control (Figure 2a). The total reads for each group were control 863, Amp 644, Neo 509, and Amp+Neo 460. The number of unique reads were control 387, Amp 200, Neo, 116, and Amp+Neo 111. Since ASV presence does not account for total reads, we next examined genera relative abundance differences between our treatment groups. Principal component analysis of the relative abundance of the top 30 identified genera resulted in separation of Amp+Neo from control (Figure 2b), but Amp+Neo did not separate from Amp or Neo along PC1 (21.4% explained variance) nor PC2 (14.67% explained variance). After determining that there are community differences between antibiotics treatments and control, we used MaAsLin2 analysis to determine which taxa of the top 30 most abundant genera were significantly associated with a specific antibiotic treatment with respect to control. This analysis revealed 18 taxa with significant associations with antibiotic treatments (Figure 2c), while the remaining 12 were not significantly associated with any antibiotic treatment. Significant associations (q < 0.05) for antibiotic-treated mice are displayed in a heatmap (Figure 2c) relative to control with their signed significance as (direction of correlation)×(− log_10_(*q*)). Thus, at a more granular level of taxonomic assignment (genera) distinct microbial communities can be associated with specific antibiotic treatments.

**Figure 2.**
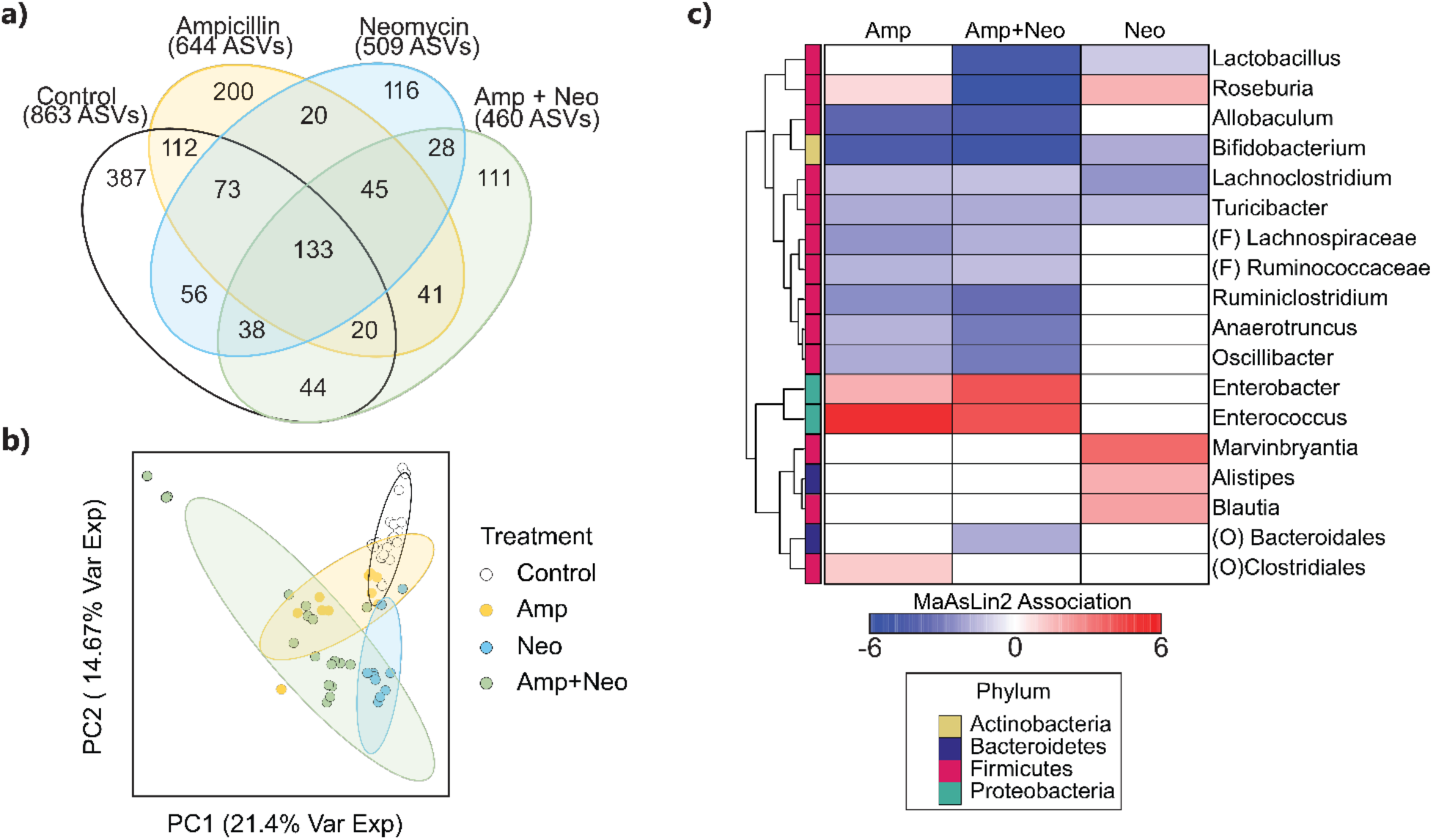
Specific taxa result in group separation among antibiotic treatments. (a) Venn diagram of unique amplicon sequence variants (ASVs) identified in control, ampicillin (Amp), neomycin (Neo), and ampicillin with neomycin (Amp+Neo) groups. (b) Principal component analysis (PCA) of genus relative abundance shows that Amp+Neo and Neo cluster away from control while Amp overlaps with control. Ellipses denote a 95% confidence interval. (c) Hierarchical clustering of significant associations of taxa relative abundance and associated phylum relative to control identified via MaAsLin2. ASVs not belonging to specific genera were assigned to their next most granular level of taxonomic separation of order (O) or family (F).

### Serum Cytokine Concentrations Are Altered after Antibiotic Treatment

Individual cytokine concentrations were measured for antibiotic treatment groups and compared to control (Supplemental Table S2). Volcano plots revealed changes in cytokine concentration for each antibiotic treatment compared to control (Figure 3a). Macrophage inflammatory protein 1 beta (MIP-1B), interleukin (IL)-6, IL-10, and macrophage colony-stimulating factor (M-CSF) concentrations are significantly lower, while IL-1a is significantly higher, with Amp. Only keratinocyte chemoattractant (KC) was significantly higher with Neo. In Amp+Neo, macrophage chemoattractant protein 1 (MCP-1) was significantly lower while IL-6 was significantly higher (Figure 3b). The significantly different circulating cytokine concentrations in each antibiotic treatment group compared to control illustrate that different antibiotic treatments result in heterogenous alteration to serum cytokine concentration.

**Figure 3.**
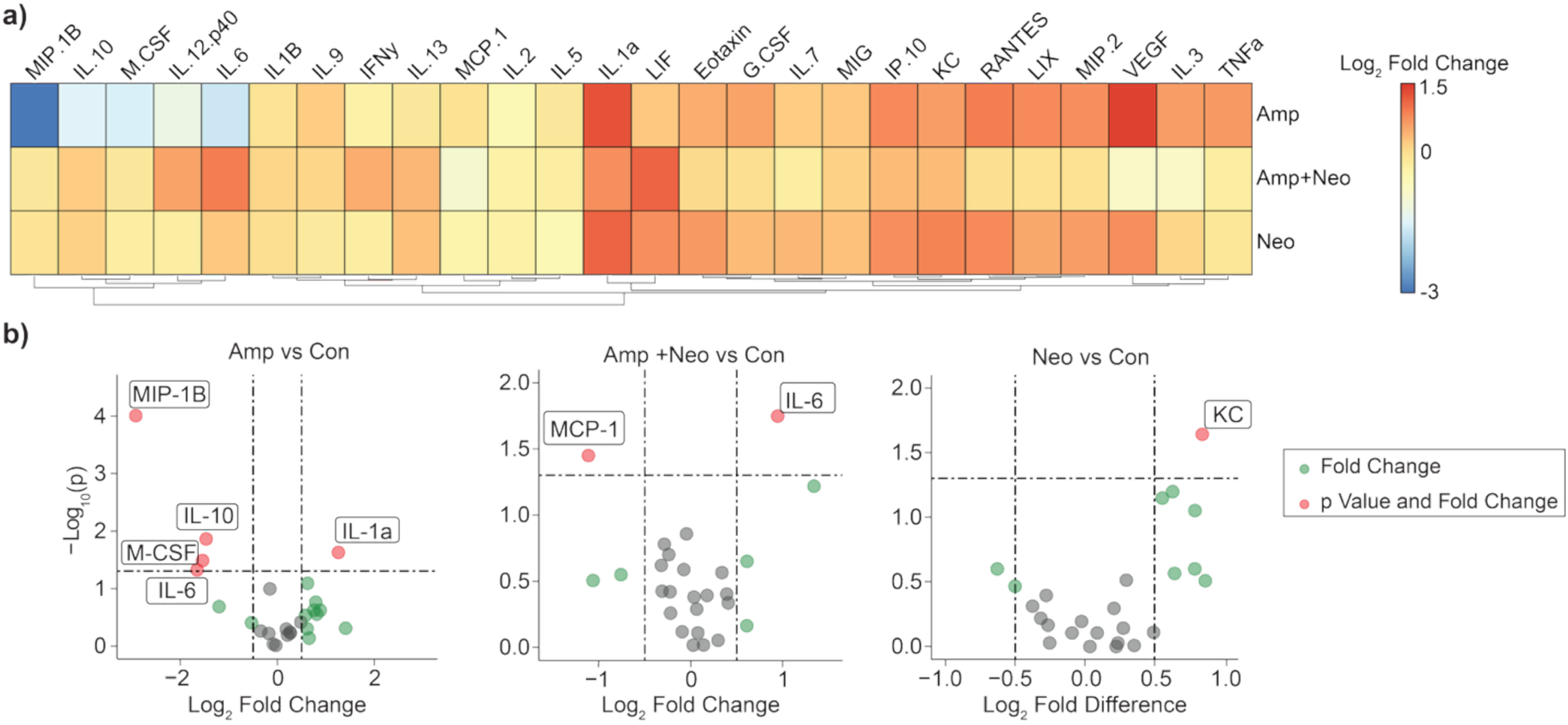
Antibiotic treatments alter serum cytokines compared to control. (a) Hierarchical clustering of log_2_ fold change of serum cytokine concentration for antibiotics treated mice relative to control. (b) Volcano plots reveal significant increase in serum cytokine concentration specific to (Con), ampicillin (Amp), neomycin (Neo), and ampicillin with neomycin (Amp+Neo) groups. Thresholds for volcano plots are p < 0.05 and log_2_(fold change) > 0.5 to capture subtle but significant changes in serum cytokine concentrations fully.

### Serum Cytokine Levels Differed among Antibiotic Treatments

Log_2_ fold change in serum cytokine concentration relative to control was significantly different in nine cytokines among the antibiotic groups. Nine cytokines were significantly altered among antibiotic treatments (Figure 4a). Eotaxin (q = 0.075), LIX (q = 0.081), RANTES (q = 0.081), and MIP-2 (q = 0.080) were significantly different among the groups and highest in Neo. RANTES (q = 0.081) and VEGF (q = 0.075) were lowest in Amp+Neo among antibiotic treatments. Levels of IL-10, IL-6, IL-13, MIP-1B, and VEGF were not different between Amp+Neo and Neo (Figure 4a). sPLS-DA on the serum cytokine concentration dataset (Figure 4b) revealed that Amp tend to cluster away from Amp+Neo and Neo along Latent Variable (LV) 1 and LV2. Serum cytokines MIP-1B, VEGF, MIP-2, IL-6, LIX, IL-10, IL-13, RANTES, Eotaxin, IL-12 (p40), M-CSF, and IP-10 had variable importance projection (VIP) scores greater than one on LV1. LV2 shared these cytokines with the addition of MCP-1. The separation of treatment groups on the derived latent variables demonstrates distinct cytokine profiles, driven by specific cytokines, between Amp+Neo and Amp, but not between Amp+Neo and Neo.

**Figure 4.**
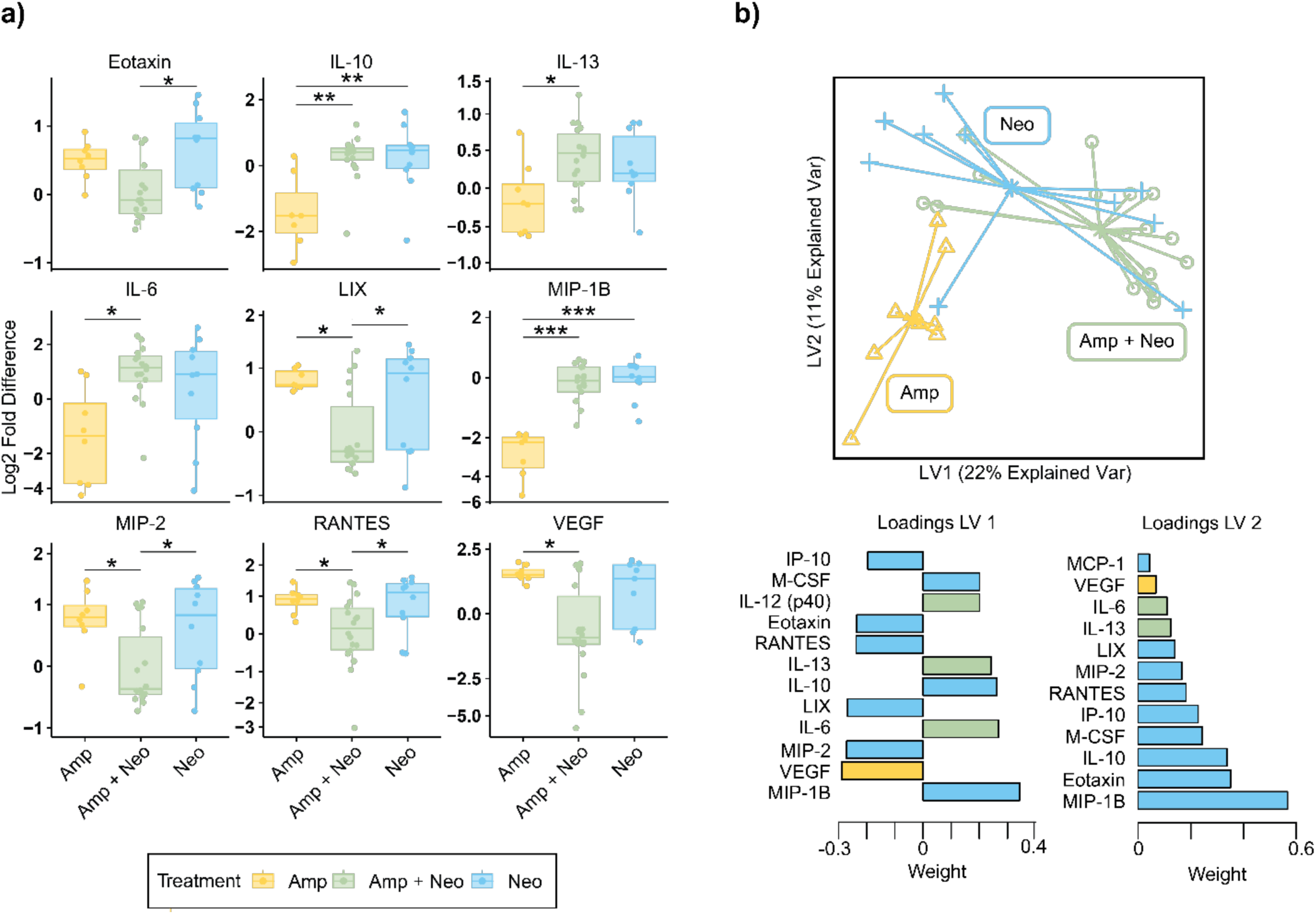
Serum cytokine concentration is altered among antibiotic treatments. (a) Log_2_(fold change) of serum cytokine concentration of cytokines with identified significant difference among ampicillin (Amp), neomycin (Neo), and ampicillin with neomycin (Amp+Neo) groups. Compared via Kruskal-Wallis corrected for false discovery rate and (*, p<0.05; **, p<0.01; ***, p<0.001) (b) Partial least squares discriminant analysis of serum cytokine fold change with variable loadings (variable importance projection, VIP >1) for latent variable 1 (LV1) and latent variable 2 (LV2) Color denotes which antibiotic treatment has the maximum weight for that cytokine.

sPLS-DA combining serum cytokine concentration and genus level relative abundance resulted in separation among Amp, Neo, and Amp+Neo (Figure 5a). Eotaxin was the only cytokine was found to have VIP > 1 on both LVs. While the taxa *Roseburia*, *Akkermansia*, Bacteroidales, Erysipelatoclostridium, Lactobacillus, Family XIII, Oscilibacter, and *Enterorhabdus* had VIP > 1 on both LV1 and LV2 (Figure 5b). Thus, including both cytokine and microbe data enables separation among antibiotic treatment groups, although it is important to note that this model more heavily weights genus relative abundance compared to cytokine concentration.

**Figure 5.**
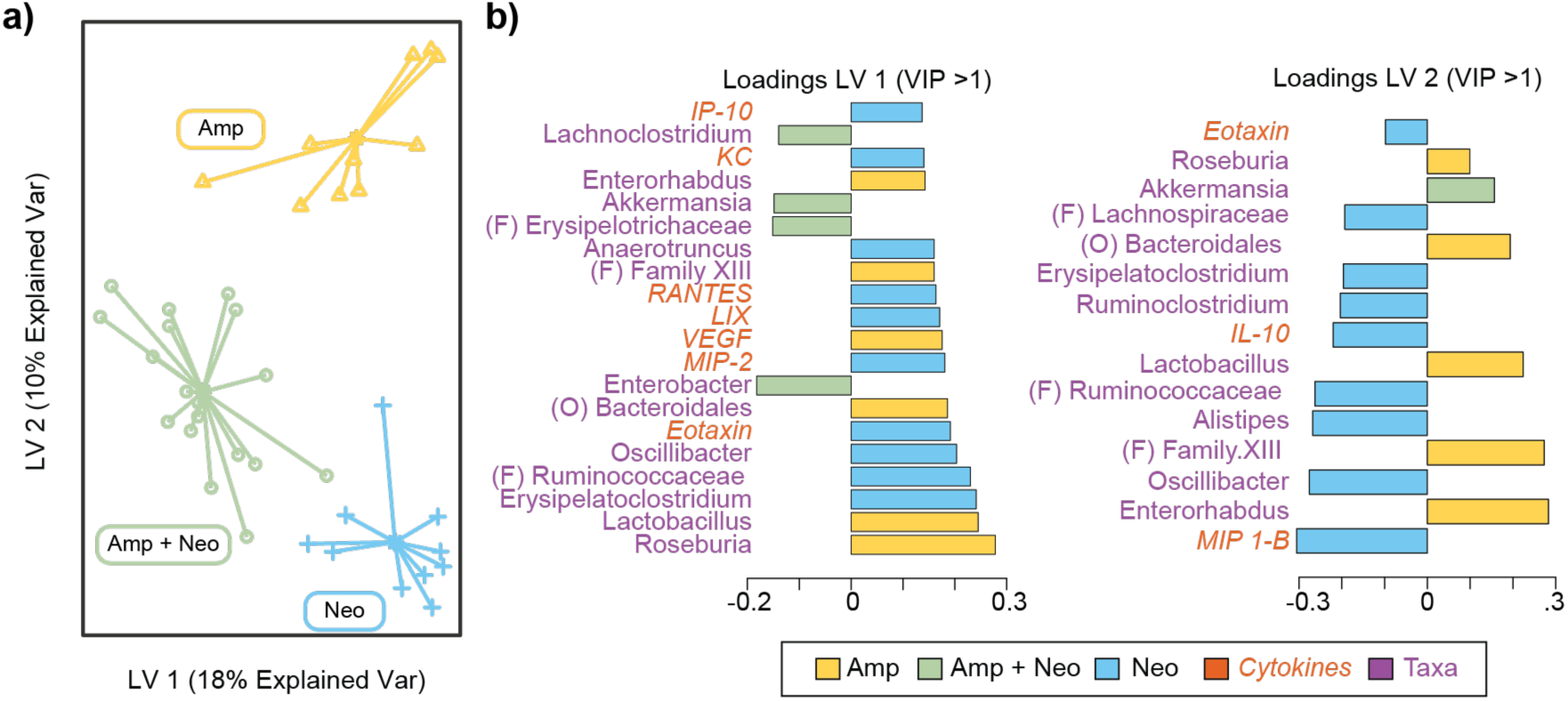
Cytokine and genera relative abundance effectively discriminates between antibiotic treatments. (a) Sparse partial least squares discriminant analysis (sPLS-DA) scores plot of ampicillin (Amp), neomycin (Neo), and ampicillin with neomycin (Amp+Neo) groups on latent variable 1 (LV1) and latent variable 2 (LV2). (b) Loadings of all variables with variable importance projection (VIP) > 1 on LV1 and LV2. Color denotes which antibiotic treatment has the maximum contribution for that cytokine.

### Treatment Duration Alters Cytokine Profile

Shannon diversity was significantly lower in short-term treatment compared to age-matched control (p <0.001, Figure 6a), but no cytokines were significantly different from control (Figure 6a). The short-term treatment also resulted in a divergent cytokine profile compared to long-term treatment (Figure 6b). sPLS-DA was able to separate the short and long treatment durations, with IL-6, Eotaxin, TNFa, IL-5, MCP-1, IL-2, and IL-1a having VIP > 1 on both LV1 and LV2 and explaining the divergent cytokine signature between short and long treatment groups (Figure 6c). These results demonstrate that downstream effects to systemic cytokines are dependent on treatment duration.

**Figure 6.**
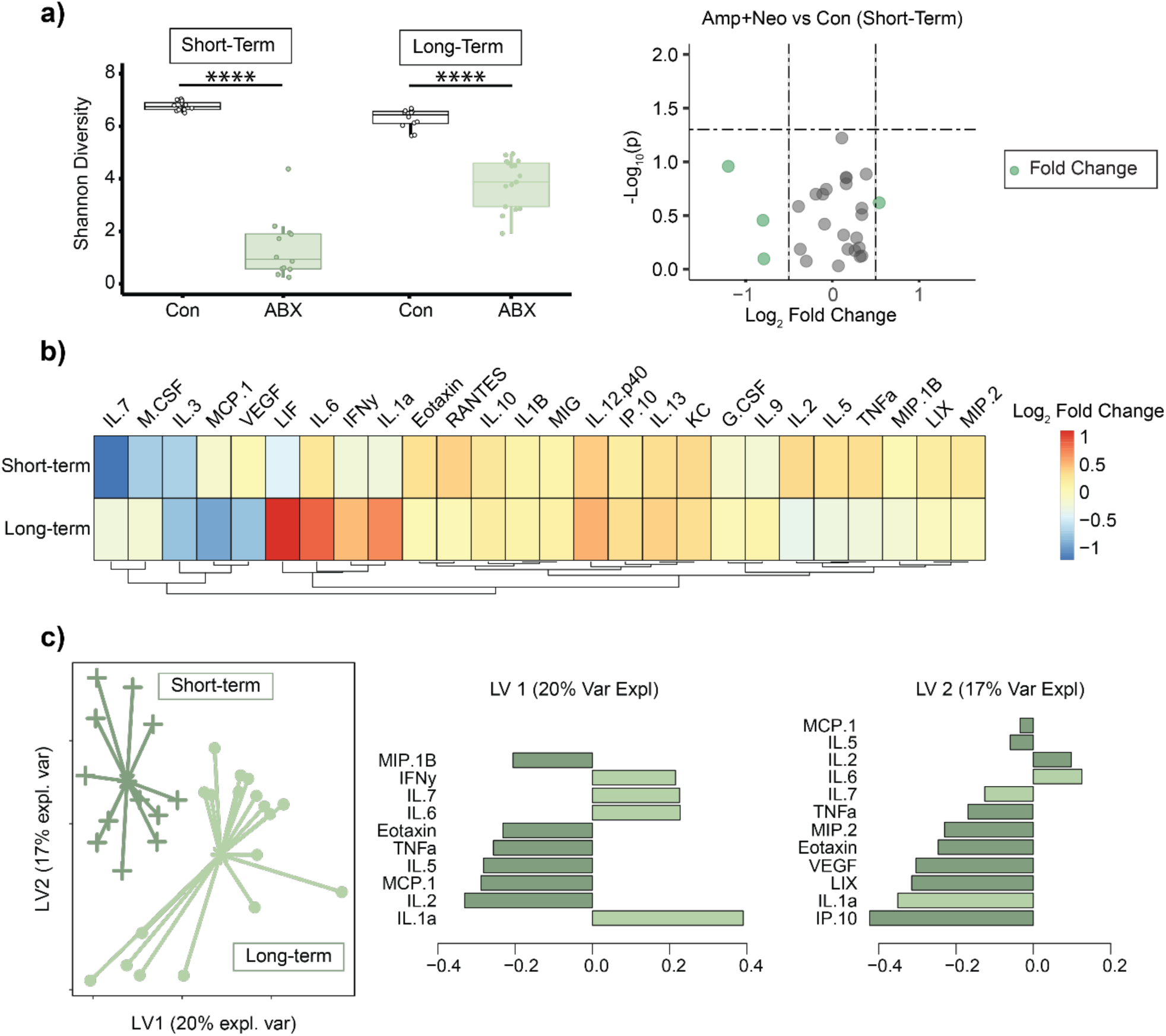
Short-term Amp+Neo treatment resulted in divergent cytokine profile compared to 12 weeks of treatment. (A) Diversity and evenness of the microbiome, as measured by Shannon diversity, was altered after ampicillin and neomycin (Amp+Neo) treatment from 5 to 8 weeks of age. However, the volcano plot of cytokine concentration for short-term Amp+Neo treatment compared to control showed no significantly altered cytokines. (B) Heatmap of fold change in cytokine concentration in mice treated in the short-vs. long-term (12 weeks, from 5 to 16 weeks of age) relative to control. (C) sPLS-DA distinguished cytokine concentrations between short- and long-term Amp+Neo treatment. Cytokines with Variable Importance Projection Score (VIP) >1 are displayed.

### Multiomic Integration and Correlation Analysis Separate Antibiotic Treatment Effects

Analysis with DIABLO revealed a correlation (r = 0.59 on LV1 and r = 0.67 on LV2) between serum cytokine concentration and genera relative abundance datasets (Figure 7a). The DIABLO model showed separation on the Genera block between all three antibiotic treatments, while only Amp separated from Neo and Amp+Neo on the cytokines block. The model was able to effectively classify treatment groups with Amp+Neo with a minimum AUC of 0.80 and highest of 0.99 (Supplemental Table S1). Correlation analysis between taxa and serum cytokines with at least one significant association with an antibiotic treatment and the 9 cytokines with significant differences among antibiotic treatment revealed specific significant correlations between taxa and cytokines. Significant correlations are displayed in a heatmap for ease of visualization (Figure 7c). Therefore, when cytokines and genera relative abundance are analyzed using multiomic integration, taxa relative abundance does not dominate the latent variable calculation, and correlative relationships between taxonomic shifts and serum cytokine concentration are readily revealed.

**Figure 7.**
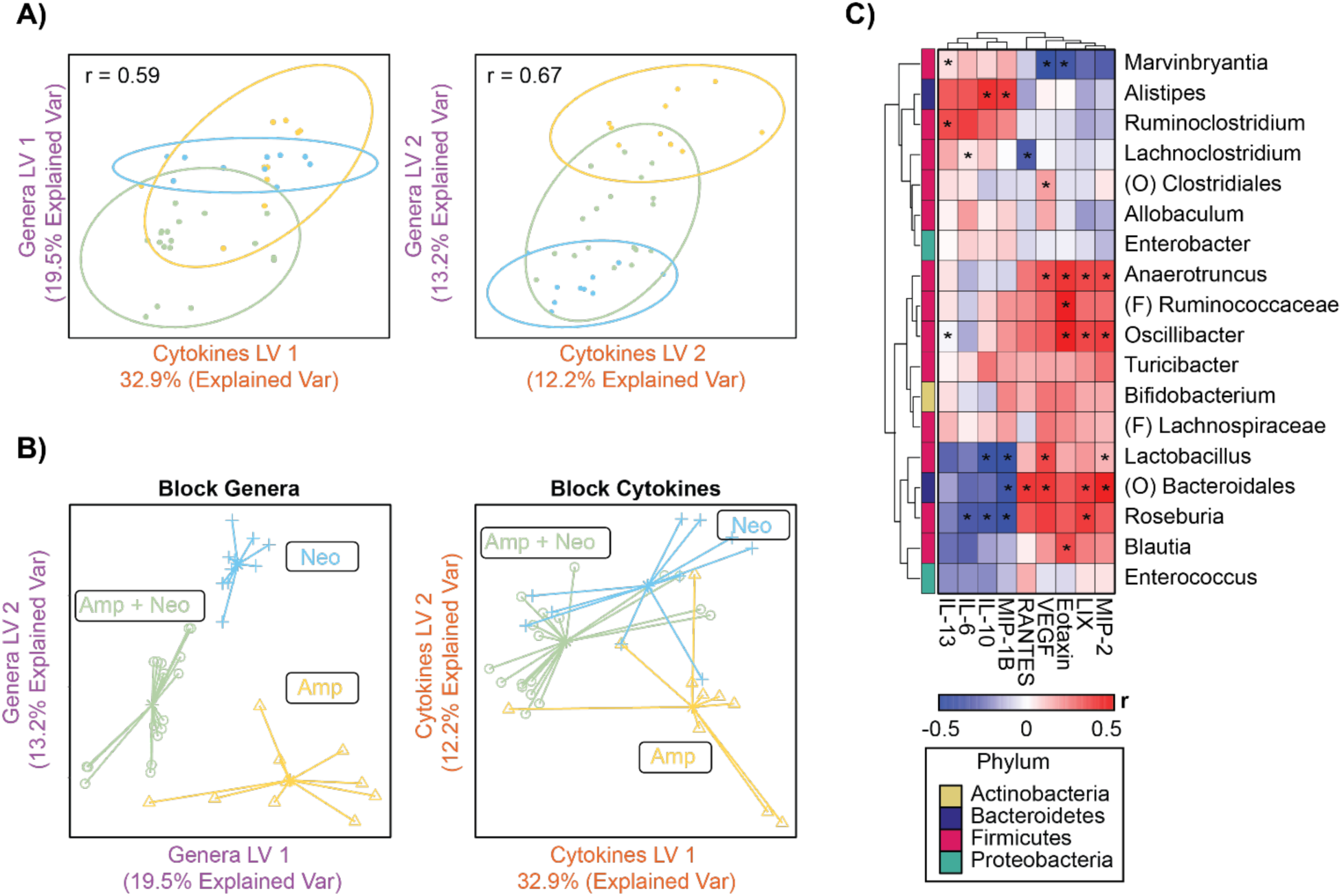
Multiomic integration reveals cytokines and taxa are significantly correlated. (A) Consensus plot derived from results of Data Integration Analysis for Biomarker discovery using Latent cOmponents (DIABLO) including ampicillin (Amp), neomycin (Neo), and ampicillin with neomycin (Amp+Neo) groups. Ellipses denote 95% confidence interval and Pearson correlation between calculated latent variables (LV) is displayed on each graph. (B) Score plots on each sparse partial least squares discriminant analysis (sPLS-DA) calculated during DIABLO. (C) Spearman correlation between serum cytokine concentration fold change and taxa. Genera are identified, except where family (F) or order (O) was the most granular identifiable taxa. (* p<0.05).

**Table 1:** Significant correlations between serum cytokine fold change and taxa relative abundance r and associated p value. Genera are identified, except where family (F) or order (O) was the most granular identifiable taxa.

| Cytokine | Taxa | $r$ | $p$ value |
| --- | --- | --- | --- |
| Eotaxin | Anaerotruncus | 0.41 | 0.01 |
|  | Blautia | 0.35 | 0.03 |
|  | (F) Ruminocaceae | 0.48 | 0.002 |
|  | Marvinbryantia | -0.4 | 0.02 |
|  | Oscilibacter | 0.48 | 0.003 |
| IL-10 | Alistipes | 0.41 | 0.01 |
|  | Lactobacillus | -0.45 | 0.01 |
|  | Roseburia | -0.4 | 0.02 |
| IL-13 | Ruminoclostridium | 0.35 | 0.04 |
| IL-6 | Roseburia | -0.37 | 0.03 |
|  | Ruminoclostridium | 0.38 | 0.03 |
| LIX | Anaerotruncus | 0.36 | 0.03 |
|  | (O) Bacteroidales | 0.39 | 0.02 |
|  | Oscilibacter | 0.37 | 0.03 |
|  | Roseburia | 0.39 | 0.2 |
| MIP-1B | Alistipes | 0.39 | 0.02 |
|  | Lactobacillus | 0.2 | <0.001 |
|  | (O) Bacteroidales | 0.39 | 0.01 |
|  | Roseburia | 0.39 | 0.01 |
| MIP-2 | Anaerotruncus | 0.36 | 0.03 |
|  | Marvinbryantia | 0.16 | 0.04 |
|  | (O) Bacteroidales | -0.43 | 0.004 |
|  | Oscilibacter | 0.21 | 0.02 |
| RANTES | Lachnoclostridium | -0.36 | 0.03 |
|  | (O) Bacteroidales | 0.4 | 0.01 |
| VEGF | Anaerotruncus | 0.34 | 0.04 |
|  | Lactobacillus | 0.37 | 0.03 |
|  | Marvinbryantia | -0.55 | <0.001 |
|  | Roseburia | 0.38 | 0.02 |

## DISCUSSION

In this work, we employed long-term antibiotics administration in mice to investigate the relationship between disruption of the gut microbiome and inflammation. We showed that the gut microbiome is altered at both the phylum and genus levels in an antibiotic-specific manner (Figures 1 and 2). We identified significant differences between the serum cytokine concentrations among treatment groups and showed that serum cytokines are altered in an antibiotic-specific manner compared to control and among treatment groups (Figure 3). Finally, through a combination of correlation (Figure 7), sPLS-DA (Figure 4), and DIABLO analysis (Figure 5), we showed that genera relative abundance and fold change of serum cytokine concentration are correlated and can discriminate between antibiotic treatments. Our results illustrate that long-term antibiotics differentially alter the gut microbiome and serum cytokine concentration and suggest potential mediators of systemic effects of microbiome disruption by ampicillin and neomycin combination treatment.

We evaluated the impact of antibiotics administration for over 2.5 months in mice, timed to skeletal maturation from 5 to 16 weeks of age. All antibiotic treatments significantly altered gut microbiome composition (Figure 1a) compared to control, illustrated by significantly lower Shannon diversity, differences in the relative abundance of several phyla, and separation from control in relative abundance of microbial genera by PCA (Figure 2b). Amp+Neo had a significantly lower relative abundance of Bacteroidetes. Some microbes in the phylum Bacteroidetes produce short-chain fatty acids (SCFAs), which have anti-inflammatory effects systemically but are reduced in inflammatory bowel disease [62]. Actinobacteria, a characteristically low abundance but broadly impactful phylum [63], was also significantly lower in Amp+Neo. Key beneficial microbes such as *Bifidobacterium* – often used in probiotic treatments due to an anti-inflammatory association [64] – belong to the Actinobacteria phylum. The relatively milder effect of individual Amp and Neo treatments on phylum relative abundance and Shannon diversity highlights that Amp+Neo promotes greater microbiome disruption and suggests potential for downstream systemic effects. Amp+Neo had the fewest unique ASVs among all groups, further illustrating the efficacy of Amp+Neo treatment in modifying the gut microbiome compared to single antibiotic treatment. Through PCA of the most abundant genera, Amp+Neo overlapped with both Amp and Neo, indicating that Amp+Neo nonetheless shares features with individual Amp or Neo treatments. Through MaAsLin, we identified 18 taxa out of the top 30 most abundant that were significantly associated with at least one antibiotic treatment (Figure 2c). Interestingly, thirteen of these taxa belong to the Firmicutes phylum, despite the relative abundance of Firmicutes having no statistical difference among any antibiotic treatment or control. Although microbes belonging to the same phylum tend to be similar, microbes within phyla may have vastly different physiological function [65]. Amp+Neo separating from control, and the many specific associations to individual taxa illustrate the dominating effects of this treatment on altering taxa, suggesting this treatment may cause the most substantial downstream changes to circulating cytokine concentration. We next sought to evaluate the systemic inflammatory state associated with these microbiome changes and found that circulating cytokine levels were altered in an antibiotic-specific manner. Among the serum cytokines significantly different with antibiotic treatment (Figure 3b), MIP-1B [66], IL-6 [67], IL-1a [68], KC [69], and MCP-1 [70] are associated with increased systemic inflammation, while IL-10 [71] and M-CSF [72] are associated with anti-inflammatory effects. We show here that Amp+Neo treatment promotes the most inflammatory state, while individual treatment of Amp or Neo alone may promote an anti-inflammatory state (Figure 3a). To identify if differences in cytokine concentration alone could discriminate among antibiotic groups we applied sPLS-DA, which effectively discriminated Amp from the other two antibiotics treatments. Neo and Amp+Neo did not separate. While neomycin is not absorbed in the gut, ampicillin is [73]. Therefore, neomycin more readily modifies gut flora than ampicillin, shown here and in other work [11, 74]. We posit that the direct effect on the gut flora by neomycin occurs in both Neo and Amp+Neo treatments, despite the lower concentration of neomycin in the combination treatment, dominating the effects of Amp.

Interestingly, we identified a generally pro-inflammatory response resulting from Amp+Neo treatment (Figure 3a), while previous studies showed an anti-inflammatory response in mice but at shorter treatment durations [25, 26]. To probe whether these differences could reflect a duration-dependent change in cytokine profile, we treated an additional set of mice from 5 to 8 weeks of age (n = 6 M/F) with Amp+Neo and compared against age-matched controls (Figure 6). We identified significant reduction to gut microbiome diversity, but no cytokines were significantly different with short-term treatment duration (Figure 6a). Compared to the longer treatment duration, short-term treatment showed a markedly less substantial change in serum cytokine concentration (Figure 6b), in agreement with studies by others [25, 26]. sPLS-DA was able to separate the short and long treatment durations. The cytokines IL-6, Eotaxin, TNFa, IL-5, MCP-1, IL-2, and IL-1a all had VIP > 1 on both LV1 and LV2 explaining the divergent cytokine signature between short and long treatment groups (Figure 6c). This additional experiment demonstrated that treatment duration affects cytokine profile and reinforced our selection of a long-term treatment duration to evaluate systemic cytokine profiles. Furthermore, this comparison also supports long-term treatment duration in future studies evaluating the role of antibiotics-induced gut disruption on processes that occur on the order of months, such as during skeletal maturation.

Since we identified significant changes to cytokine profile only at long-term treatment duration, we focused multiomic methods on this time point. Noting that the antibiotic treatments clustered differently in microbiome-only compared to cytokine-only analyses, we combined the microbiome and cytokine datasets to determine if together they would better discriminate between antibiotic treatments via sPLS-DA. All three treatment groups formed distinct clusters. On LV1, *Lactobacillus* and *Roseburia* had the largest loadings with VIP > 1 of all taxa and discriminated for Amp. Notably, *Lactobacillus* and *Roseburia* were also significantly associated with a specific treatment via MaAsLin and are considered beneficial and impactful taxa in the gut microbiome [75, 76]. sPLS-DA effectively discriminated between antibiotic treatment groups, and the model skewed toward microbiome relative abundance. Of all variables with VIP > 1, 13 taxa and 7 cytokines were found in LV 1, and 12 taxa and 3 cytokines were found in LV2. The greater weight given to taxonomic data on each LV illustrates the need for multiomic integration to enable a more balanced analysis of how microbiome composition may drive different serum cytokine levels.

Using DIABLO for multiomic integration, we identified moderate correlations between genera relative abundance and serum cytokine concentration on LV1 (r = 0.59) and LV2 (r = 0.67), demonstrating the interrelatedness of inflammation and microbiome manipulation (Figure 7). Through this integrative approach, Amp and Neo separated on the consensus plot of LV1 from the DIABLO block sPLS-DA (Figure 7b). We compared this analysis with Spearman correlation to identify specific associations between cytokine concentrations and taxa. Notably, IL-6, IL-10 and MIP-1B were all significantly negatively correlated with *Roseburia*, a microbe typically associated with anti-inflammatory properties [77, 78]. *Roseburia* treatment reduced colitis associated intestinal inflammation [79] and *Roseburia*’s relative abundance is lower in rheumatoid arthritis patients [80]. IL-10 and MIP-1B were both significantly negatively correlated with *Lactobacillus*, which is largely considered a beneficial microbe. Treatment with *Lactobacillus* protected against pathogenic intestinal barrier disruption *in vitro* [81] and *Lactobacillus* species reduce inflammatory cytokine levels *in vitro* [82]. Furthermore, *Lactobacillus* protects against bone loss in ovariectomized mice [83] and mitigates bone loss in older women [84]. IL-10 deficient mice have increased intestinal permeability [85]. Mucosal and serum concentration of IL-6 are both increased with inflammatory bowel disease in humans and animal models [86–88]. MIP-1B stimulates mucosal immune function [89]. Among the various correlations between microbial taxa and cytokines (Figure 8), IL-6, MIP-1B, and IL-10 levels were significantly altered with antibiotics consistently over multiple levels of analysis, suggesting the greatest sensitivity to microbiome manipulation. These cytokines were significantly up- or down-regulated compared to control, were significantly different among antibiotic treatments, had VIP > 1 on the cytokine-specific sPLS-DA, and showed a significant correlation with at least one microbial taxon. Notably, these cytokines are also associated with synovial joint health and disease (Figure 8, Supplemental Table S3) [31, 32, 34, 36, 89–113]. Osteoarthritic synovial fluid has increased concentration of MIP-1B [114]. IL-10 protects against cartilage degradation *in vitro* [115], and IL-10 deficient mice experience bone loss and frailty [96]. IL-6 increases glycosaminoglycan degradation in cartilage [94] and promotes pathologic fibroblast activity in tendinopathy [37, 38]. IL-6 has been shown to promote bone formation [116], but it is also shown to drive osteoclast differentiation and joint degradation in rheumatoid arthritis [117]. The multifaceted roles of these cytokines in both the gut and synovial joints and associations with microbial taxa illustrate their potential to mediate putative gut-joint interactions and are worthy of future investigation.

**Figure 8.**
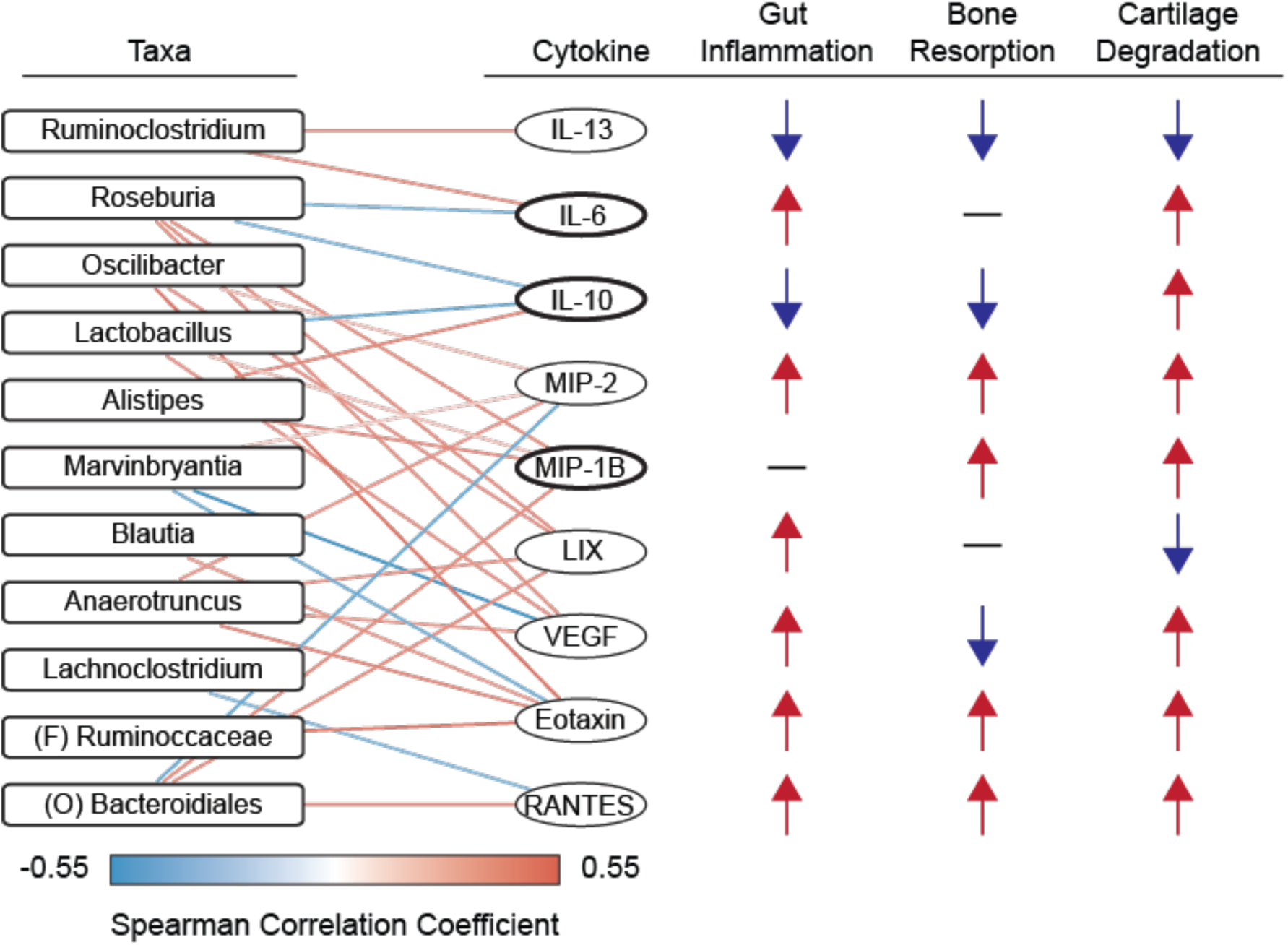
Cytokines with significant correlation to taxa relative abundance are also associated with gut and joint physiologic and pathophysiologic processes. Cytokines with significant spearman correlation to at least one taxon are displayed on the left, and their association with gut inflammation, bone resorption, and cartilage degradation identified through literature are shown on the right. An up arrow (red) indicates that the cytokine has a positive association with that process, while a down arrow (blue) indicates that the cytokine has a negative association with that process. The cytokines IL-6, IL-10, and MIP-1B (bolded ellipses) were consistently different across multiple levels of analysis indicating that these cytokines are sensitive to microbiome manipulation and may be promising targets for continuing analysis. Citations for relevant literature are displayed in Supplemental Table S3.

In conclusion, we have demonstrated antibiotic-specific changes to serum cytokine concentration and gut microbiome community and shown through sPLS-DA and multiomic integration that serum cytokines and 16S metagenomics, considered together, enable discrimination between antibiotic treatments. We also showed antibiotic-specific associations among specific taxa and serum cytokines. Notably, we identified three key cytokines (IL-6, IL-10, MIP-1B) and known mediators of both synovial joint and intestinal health and disease were consistently perturbed with antibiotic treatments. Because bone deficits have previously been associated with ampicillin and neomycin cocktail [9, 10, 48, 118], which we now show to be pro-inflammatory, this work also represents an initial step towards studying the mechanisms connecting the gut-host interaction with synovial joint health and disease – the “gut-joint axis”. Characterizing the associations between antibiotic-specific alteration of the gut microbiome and serum cytokine concentration thus improves our understanding of how microbiome modulation may in turn affect synovial joint health and disease, providing a foundation for future gut-joint axis studies.

## ACKNOWLEDGEMENT

We acknowledge help from Zachary R. Davis on specimen preparation.

## CITATION DIVERSITY STATEMENT

The scientific literature has documented biases in citation practices, including under-citation of minority scholars. We recognize this bias and have worked to ensure that we are referencing appropriate papers with fair author inclusion

## ETHICAL APPROVAL

All *in vivo* murine experiments were conducted under institutional approval (Purdue IACUC protocol 2104002138). C57Bl6/J mice were bred in conventional housing in AALAAC-compliant facilities.

## CONSENT TO PARTICIPATE

Not applicable

## CONSENT TO PUBLISH

Not applicable

## DATA AVAILABILITY STATEMENT

Collected data is made available through a Harvard Dataverse data repository at https://doi.org/10.7910/DVN/ZQ2FXS.

## AUTHORS CONTRIBUTIONS

Both authors contributed to the study conception and design. Formal data collection and analysis were performed by CXV. All authors wrote, reviewed, and edited the manuscript. Both authors read and approved the final manuscript.

## FUNDING

This work was supported in part by a Defense Advanced Research Projects Agency Young Faculty Award, funded through the Army Research Office as Contract W911NF2110372.

## COMPETING INETERESTS

The authors have no competing interests to disclose.

## Notes

https://doi.org/10.7910/DVN/ZQ2FXS

### Competing Interest Statement

The authors have declared no competing interest.

### Summary of Updates

Updated text to reflect additional analysis of a short-term (3-4 week) treatment with ampicillin and neomycin in comparison to the long-term treatment with the same antibiotic cocktail. Results show that short- and long-term treatment result in a differing cytokine profile in mice that are treated starting at 5 weeks of age. Additionally, figures are adjusted to have more consistently formatting, typographic errors are fixed, and an additional discussion is complemented with a figure that summarizes the microbe to cytokine associations measured in this study as well as how those cytokines have been linked in the literature to gut and joint changes.

