## Supplemental Table S1 for "Multiomic Integration Reveals Taxonomic Shifts Correlate to Serum Cytokines in an Antibiotics Model of Gut Microbiome Disruption"

***Table S1: Multiomic integration effectively classified antibiotic treatments.*** *Area under the curve (AUC) for Data Integration Analysis for Biomarker discovery using Latent cOmponents (DIABLO) and associated p-value determined using mixOmics R package.*

| Latent Variable | Block | Comparison | AUC | P value |
|---|---|---|---|---|
| LV 1 | Genera | Amp vs. others | 0.80 | 0.004 |
| | | Neo vs. others | 0.81 | 0.004 |
| | | Amp+Neo vs. others | 0.98 | <0.001 |
| | Cytokines | Amp vs. others | 0.66 | 0.14 |
| | | Neo vs. others | 0.74 | 0.03 |
| | | Amp+Neo vs. others | 0.81 | 0.001 |
| LV 2 | Genera | Amp vs. others | 0.95 | <0.001 |
| | | Neo vs. others | 0.89 | <0.001 |
| | | Amp+Neo vs. others | 0.99 | <0.001 |
| | Cytokines | Amp vs. others | 0.90 | <0.001 |
| | | Neo vs. others | 0.81 | 0.003 |
| | | Amp+Neo vs. others | 0.80 | 0.001 |
