## Supplemental Table S2 for "Multiomic Integration Reveals Taxonomic Shifts Correlate to Serum Cytokines in an Antibiotics Model of Gut Microbiome Disruption"

***Table S2: Serum cytokine concentrations are different with respect to control.*** *Serum cytokine concentrations reported as mean ± SEM. Bold text denotes a statistically significant difference ($p < 0.05$) compared to control. Tested with Kruskal Wallis and subsequent Dunn testing with Benjamini-Hochberg correction.*

| Cytokine | Control | Amp | Amp+Neo | Neo |
|---|---|---|---|---|
| Eotaxin | 2182.3 ± 264.9 | 2656.9 ± 35 | 2039.3 ± 157.8 | **3104.4** ± **373.9** |
| Granulocyte-colony stimulating factor (G-CSF) | 572.3 ± 402.9 | 708.1 ± 344.8 | 422.4 ± 137.7 | 624.2 ± 462.8 |
| Interferon gamma (IFNγ) | 12.4 ± 8.5 | 10.0 ± 6.8 | 16.4 ± 13.5 | 8.4 ± 3.6 |
| Interleukin (IL)-10 | 45.4 ± 25.3 | 17.2 ± 14.2 | 43.0 ± 14.4 | 48.8 ± 25.7 |
| IL-12(p40) | 70.3 ± 56.5 | 38.1 ± 39.1 | 73.1 ± 40.6 | **56.6** ± **55.9** |
| IL-13 | 136 ± 59.5 | 109.5 ± 40 | 158.1 ± 47.3 | 142.9 ± 46.1 |
| IL-1β | 7.2 ± 4.6 | 7.4 ± 5 | 8.0 ± 3.9 | 7.4 ± 4.3 |
| IL-1a | 356.3 ± 337.3 | 618.2 ± 249.1 | 437.1 ± 286.4 | **822.7** ± **768.9** |
| IL-2 | 10.6 ± 6.6 | 7.5 ± 4.7 | 7.6 ± 3.4 | 8.1 ± 4.1 |
| IL-3 | 5.4 ± 4.8 | 7.8 ± 7.5 | 3.0 ± 2.9 | 4.9 ± 4.2 |
| IL-5 | 33.6 ± 17.9 | 39.7 ± 38.8 | 27.9 ± 17.7 | 26.9 ± 14.0 |
| IL-6 | 12.7 ± 7.2 | 6.6 ± 7.2 | **23.1** ± **13.0** | 21.5 ± 21.3 |
| IL-7 | 6.7 ± 4.4 | 5.7 ± 2.7 | 4.5 ± 2.6 | 7.3 ± 4.3 |
| IL-9 | 217.7 ± 112.9 | 294.2 ± 202.6 | 227.5 ± 109.0 | 202.5 ± 135.0 |
| Interferon-gamma induced protein (IP-10) | 411.0 ± 266.4 | 559.2 ± 125.8 | 418.3 ± 170.4 | **602.5** ± **254.8** |
| Keratinocyte chemoattractant (KC) | 117.8 ± 72.0 | 161.4 ± 54.6 | 131.8 ± 43.5 | **186.2** ± **60** |
| Leukemia Inhibitory Factor (LIF) | 11.0 ± 8.7 | 8.6 ± 4.6 | **19.7** ± **20.7** | 9.8 ± 5.3 |
| Lipopolysaccharide-induced CXC chemokine (LIX) | 3161.1 ± 1894.2 | **4514.6** ± **528.4** | 2705.9 ± 1451.4 | **4246.5** ± **1935.1** |
| Macrophage colony-stimulating factor (M-CSF) | 42.0 ± 43.7 | 13.4 ± 12.8 | 27.9 ± 21.3 | 30.5 ± 21.8 |
| Macrophage chemoattractant protein (MCP-1) | 85.9 ± 86.8 | 64.2 ± 36.4 | 37.4 ± 34.4 | 48.4 ± 33.2 |
| Monokine induced by gamma (MIG) | 1274.6 ± 861.7 | 1311.4 ± 517.2 | 1244 ± 669.1 | 1365.4 ± 87.9 |
| Macrophage inflammatory protein (MIP-1β) | 64.3 ± 45.6 | **8.2** ± **4.8** | 43.6 ± 15.3 | 49.7 ± 16.6 |
| MIP-2 | 481.7 ± 309.3 | 629.6 ± 203.1 | 374.7 ± 188 | **620.4** ± **283.0** |
| Regulated on Activation, Normal T Expressed and Secreted (RANTES) | 56.9 ± 49.7 | **70.5** ± **16.9** | 45.2 ± 25.6 | 71.3 ± 30.0 |
| Tumor Necrosis Factor alpha (TNFα) | 13.2 ± 11.6 | 15.2 ± 9.5 | 7.8 ± 5.9 | 8.2 ± 4.2 |
| Vascular endothelial growth factor (VEGF) | 6.9 ± 5.9 | 9.9 ± 2.1 | 4.2 ± 4.4 | **8.3** ± **4.8** |
