## Supplemental Table S3 for "Multiomic Integration Reveals Taxonomic Shifts Correlate to Serum Cytokines in an Antibiotics Model of Gut Microbiome Disruption"

***Table S3: Cytokines with significant correlation to taxa relative abundance are also associated with gut and joint physiologic and pathophysiologic processes.***

| **Cytokine** | **Gut Inflammation** | **Bone Restitution** | **Cartilage Degradation** |
|---|---|---|---|
| IL-13 | Donlan, *et al.*, 2024 [90] | Onoe, *et al.*, 1996 [91] | Nabbe, *et al.*, 2005 [92] |
| IL-6 | Shahini, *et al.*, 2023 [93] | Yoshitake, *et al.*, 2008 [32]<br>Ishimi, *et al.*, 1990 [31] | Flannery, *et al.*, 2000 [94] |
| IL-10 | Kuhn, *et al.*, 1993 [95] | Dresner-Pollak, *et al.*, 2004 [96] | Behrendt, *et al.*, 2018 [97]<br>Kasama, *et al.*, 1995 [98] |
| MIP-2 | Ohtsuka, *et al.*, 2001 [99] | Ha, *et al.*, 2011 [100] | Kasama, *et al.*, 1995 [98] |
| MIP-1B | Lillard, *et al.*, 2003 [89]<br>Grimm, *et al.*, 1996 [101] | Abe, *et al.*, 2002 [34] | Stucker, *et al.*, 2025 [36] |
| LIX | Kwon, *et al.*, 2005 [102] | Klosterhoff, *et al.*, 2022 [103]<br>Sundaram 2013 [104] | Kawata, *et al.*, 2021 [105] |
| VEGF | Scaldaferri, *et al.*, 2009 [106] | Hu, *et al.*, 2016 [107] | Nagao, *et al.*, 2017 [108] |
| Eotaxin | Coburn, *et al.*, 2013 [109] | Ahmadi, *et al.*, 2020 [110] | Hsu, *et al.*, 2004 [111] |
| RANTES | Grimm, *et al.*, 1996 [101] | Lechner, *et al.*, 2018 [112] | Alaaeddine, *et al.*, 2001 [113] |
